# Automated inference of disease mechanisms in patient-hiPSC-derived neuronal networks

**DOI:** 10.1101/2024.05.23.595522

**Authors:** Nina Doorn, Michel J.A.M. van Putten, Monica Frega

**Affiliations:** Department of Clinical Neurophysiology, University of Twente, 7522 NB Enschede, The Netherlands; Department of Neurology and Clinical Neurophysiology, Medisch Spectrum Twente, 7512 KZ Enschede, The Netherlands

## Abstract

Human induced pluripotent stem cells (hiPSCs)-derived neurons offer a valuable platform for studying neurological disorders in a patient-specific manner. These neurons can be rapidly differentiated into excitatory neuronal networks, whose activity is measurable using multi-electrode arrays (MEAs). Neuronal networks derived from patients exhibit distinct characteristics, reflecting underlying pathological molecular mechanisms. However, elucidating these mechanisms traditionally requires extensive and hypothesis-driven additional experiments. Computational models can link observable network activity to underlying molecular mechanisms by identifying biophysical model parameters that simulate the activity, but this identification process is challenging. Here, we address this challenge by using simulation-based inference (SBI), a machine-learning approach, to automatically identify the full range of model parameters able to explain the patient-derived MEA activity.

Our study demonstrates that SBI can accurately identify ground-truth parameters in simulated data, and successfully estimate the parameters that replicate the network activity of healthy hiPSC-derived neuronal networks. Furthermore, we show that SBI can pinpoint molecular mechanisms affected by pharmacological agents and identify key disease mechanisms in neuronal networks derived from patients. These findings underscore the potential of SBI to automate and enhance the identification of disease mechanisms from MEA measurements, offering a robust and scalable method for advancing research with hiPSC-derived neuronal networks.

## Introduction

Human induced pluripotent stem cells (hiPSCs)-derived neurons have become a key technology to investigate neurological disorders using a patient-specific background in a controlled in vitro environment. Specifically, by differentiating hiPSCs into excitatory neurons through forced expression of *Ngn2*, researchers can rapidly generate electrically mature neuronal networks, whose activity can be easily measured using multi-electrode arrays (MEAs) [1]. *In vitro* neuronal networks derived from healthy subjects or patients show robust and replicable functional phenotypes on MEAs [2], and various genotype/phenotype correlations have been established using this platform [3, 4, 5, 6, 7]. In this way, MEA data from patient-derived neuronal networks can be obtained efficiently with high throughput. Since the electrical activity of the neuronal networks is shaped by underlying molecular mechanisms, the MEA data may contain valuable information about these underlying processes [8]. However, unveiling these mechanisms requires additional extensive and hypothesis-driven *in vitro* experiments, which are both time-consuming and costly. This is further complicated by the many interacting mechanisms that influence network activity, and the possibility that similar activity patterns can arise from different underlying properties.

Computational models are invaluable in bridging the gap between experimental observations and the (patho-)physiological mechanisms underlying them [9]. Previously, we have developed a bio-physically detailed computational model of hiPSC-derived excitatory neuronal networks on MEA [10, 11], that can dissect the effect of specific cellular changes on network activity. The parameters of this *in silico* model describe key physiological characteristics of the neurons, the synapses, and the network. By adjusting these parameters, simulations that faithfully resemble either healthy-or patient-derived neuronal network activity can be obtained. Differences between the parameters of healthy and disease models reflect altered biological properties in the diseased network. However, finding the optimal model parameters able to reproduce specific experimental data is difficult.

The optimal parameters cannot be calculated explicitly in most biophysical models, because the models are only defined implicitly with stochastic computer simulations. Often, trial-and-error is used to find suitable parameters, but this is a time-consuming and not systematic approach. An alternative is performing many simulations with different combinations of parameters drawn from a grid or following a specific route through parameter space, and evaluating which simulation matches the experimental observation best [12, 13]. These and similar methods have several limitations. First, the user is required to define a distance measure describing how well the simulation replicates the experimental measurement, which can be challenging when observations have many features. Second, these methods quickly become computationally expensive when more parameters need to be identified and the amount of required simulations exponentially increases, especially since these simulations need to be repeated for every experimental observation. Third, most methods provide a single “best” set of parameters, without information about other possible parameter combinations or the uncertainty of the estimation. This is especially a limiting factor in neuronal modeling where different parameter combinations can result in similar observations (parameter degeneracy) [14], and where activity is naturally robust to some parameter perturbations but very sensitive to changes in other parameters [15].

To address these limitations, simulation-based inference (SBI) was introduced [16]. SBI is a machine-learning approach that allows efficient statistical inference of biophysical model parameters using simulations, only. SBI can identify not only the best parameters, but the full set of parameter combinations for which the model reproduces the experimental measurements, and their corresponding probabilities (i.e., the posterior distribution). Moreover, the process is amortized, meaning heavy computations only need to be performed once, after which new experimental measurements can be readily analyzed.

Here, we apply SBI to our previously validated computational model of hiPSC-derived neuronal networks on MEA [10]. Our primary aim is to assess the suitability of SBI for the automatic identification of disease mechanisms underlying the phenotype of patient-derived neuronal networks. We obtained the parameters’ posterior distribution using multiple experimental MEA measurements from healthy-, and patient-derived neuronal networks. This was complemented by experimental data from neuronal cultures treated with different pharmacological agents. First, we show that the most probable parameters identified by SBI result in simulations with high similarity to experimental measurements. Second, we show that SBI identifies phenotype-causing molecular mechanisms that have previously been proven to be affected in patient-derived networks in vitro. Finally, we provide a set of recommendations for the use of SBI to identify disease mechanisms from MEA measurements.

## Results

### SBI correctly identifies ground-truth parameters of simulated neuronal-network activity

We used SBI to infer mechanistic information from measurements of hiPSC-derived excitatory neuronal networks on MEA (Figure 1A). To do so, we sampled 300,000 model parameter configurations from a prior distribution (i.e., a uniform distribution within plausible parameter ranges, see Table 1) (Figure 1A1) and used them for simulations with our biophysical computational model (Figure 1A2). We analyzed the resulting simulations by computing 13 MEA features that capture the most important characteristics of the network activity and are often used in MEA literature (see Methods and Table 2 for details). We then used the analyzed simulations to train a deep neural density estimator (NDE) to identify which model parameter sets produce simulations compatible with experimental measurements (Figure 1A3). We then evaluated the trained NDE with the MEA features of recordings from healthy and diseased hiPSC-derived neuronal networks (Figure 1A4, A5). This resulted in the full space of model parameters compatible with the experimental observations (Figure 1A6). This space is called the posterior distribution and shows a high probability (yellow) for model parameters consistent with the data and a probability close to zero (dark blue) for model parameters not leading to data mimicking simulations (Figure 1A7).

**Table 1:**
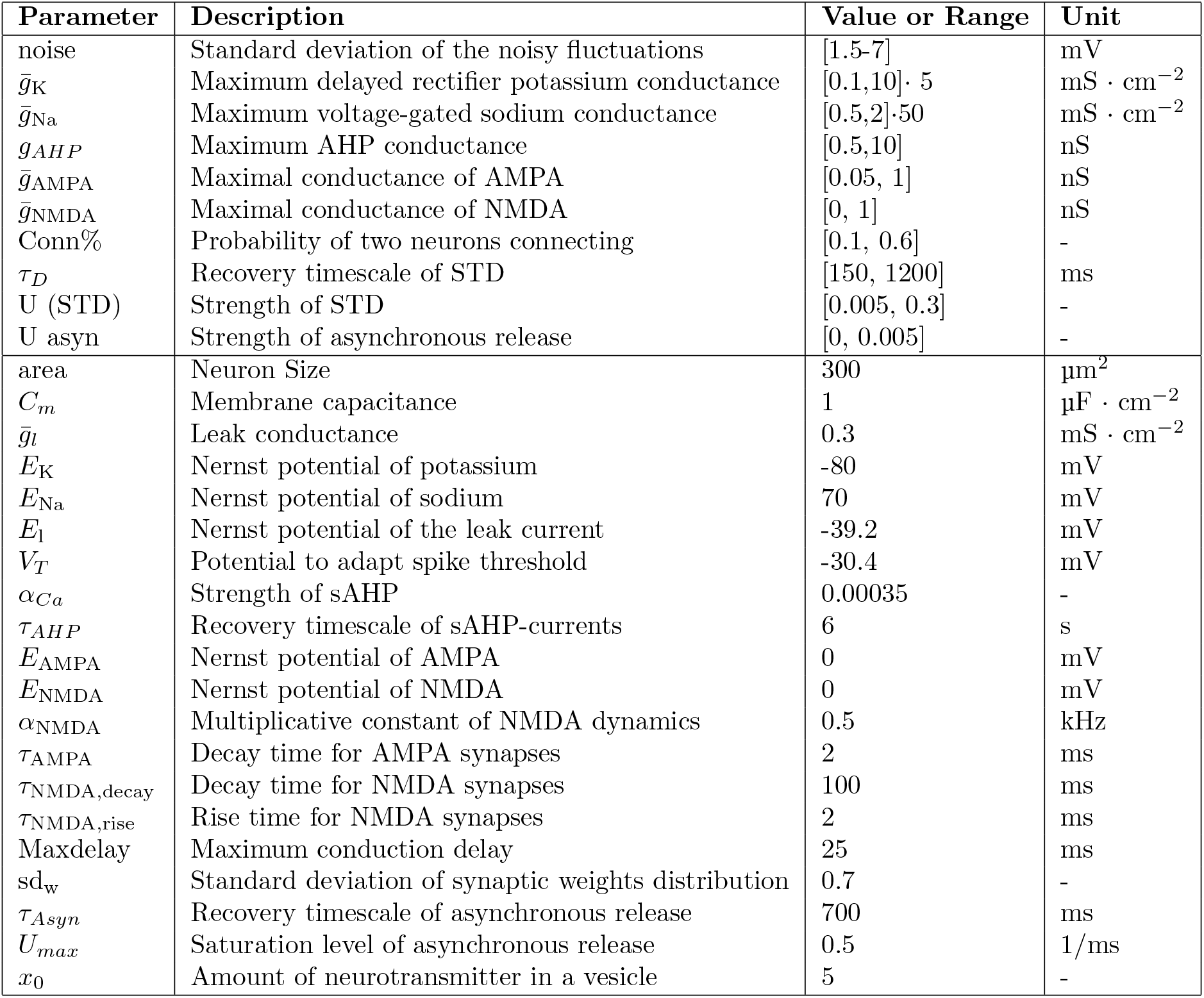
Overview of the parameter value ranges of the free parameters, and values of the fixed parameters of the computational model.

**Table 2:** Overview of the MEA features used for training of the NDE.

| MEA feature | Description | Unit |
| --- | --- | --- |
| MFR | Mean Firing Rate | Hz |
| NBR | Network Burst Rate | 1/min |
| NBD | Network Burst Duration | s |
| PSIB | Percentage of Spikes in NBs | - |
| #FBs | Average number of fragments per NB | - |
| CV<sub>IBI</sub> | Coefficient of Variation of the Inter-NB-Intervals | - |
| mean ISI CC | Average Pearson Correlation Coefficient between ISI-timeseries | - |
| sd ISI CC | Standard deviation of the Pearson Correlation Coefficients | - |
| ISI dist | Average ISI distance between ISI-timeseries of electrode-pairs | - |
| mean ISI | The average ISI between all electrodes and timepoints | s |
| sd ISI temp | Standard deviation of ISI-values averaged between electrodes | s |
| sd ISI elec | Standard deviation of ISI-values averaged over time | s |
| MAC | Maximum Autocorrelation Component | - |

**Figure 1:**
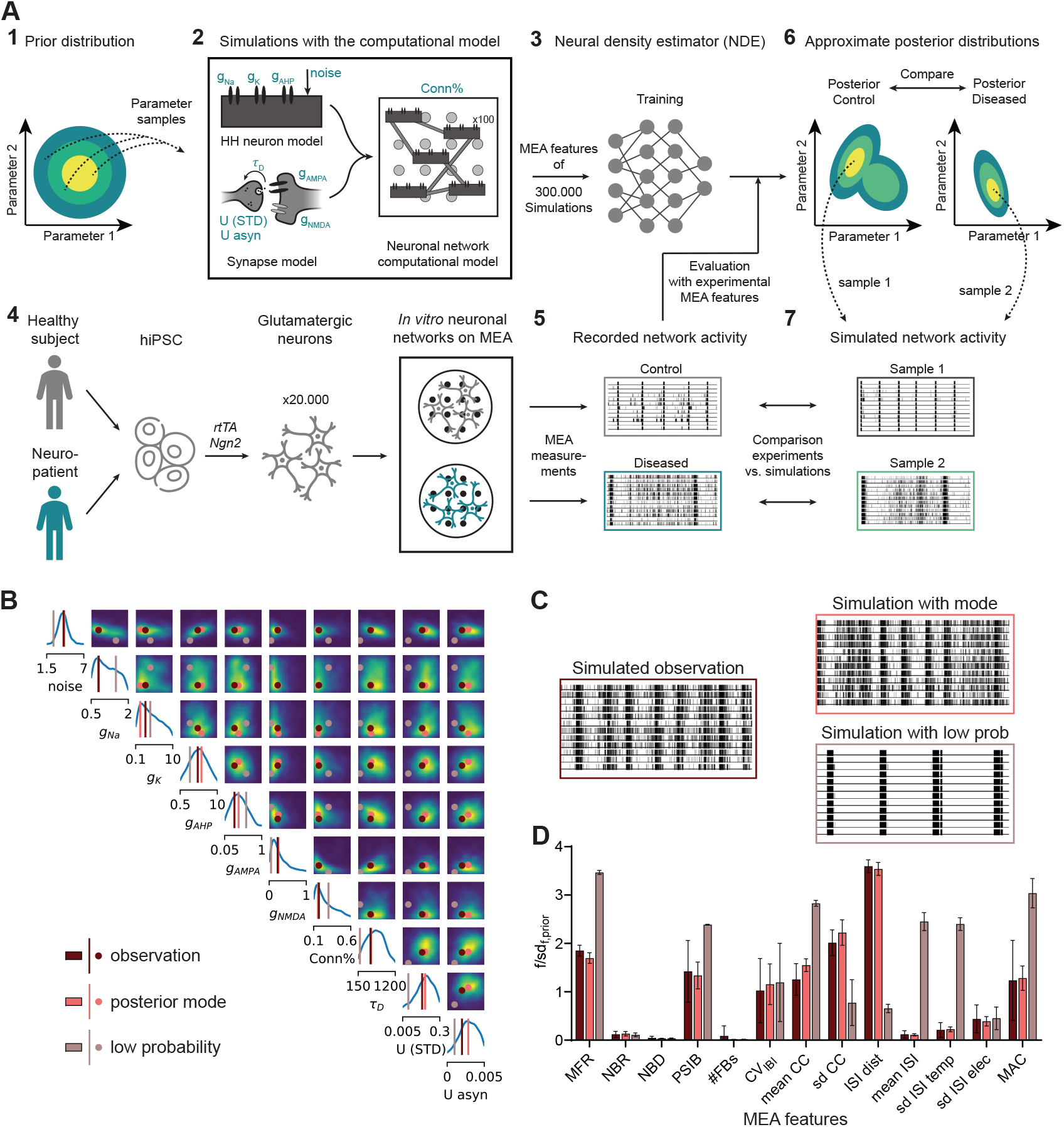
SBI to infer the entire space of parameters consistent with experimental MEA measurements. **A)** Schematic overview of the method: **1)** 300,000 parameter configurations are drawn from the prior distribution; **2)** simulations are performed using our biophysical computational model consisting of 100 Hodgkin-Huxley (HH)-type neurons and AMPAr, and NMDAr-mediated synapses. Model parameters are depicted in blue and reported in Table 1; **3)** MEA features (reported in Table 2) of the resulting 300,000 simulations are used to train a neural density estimator (NDE); **4)** schematic overview of the protocol to differentiate hiPSCs into neuronal networks on multi-electrode arrays (MEAs): hiPSCs were obtained by reprogramming somatic cells of healthy subjects and patients, excitatory neurons were generated through doxycycline (Dox) inducible overexpression of Neurogenin2 (*Ngn2*), activity was recorded around DIV 35 with MEA; **5)** MEA features are computed from the recorded network activity and used to evaluate the trained NDE; **6)** this resulted in the posterior distribution per experimental observation, which can be compared to identify mechanistic differences; **7)** simulations with parameters sampled from the posterior distribution are similar when compared to experimental observations. **B)** Inferred posterior of the ten model parameters given 13 MEA features of simulated activity. Diagonal shows univariate marginals and off-diagonal shows pairwise marginals between the two adjacent parameters, where a lighter color indicates a higher probability. Ground-truth model parameters are shown in brown, posterior mode in pink, and low-probability model parameters in beige. **C)** Rasterplots showing 1 minute of (left) simulation used as input for the inference, (top right) example simulation with the mode of the posterior distribution, and (bottom right) example simulation with the low probability model parameters. **D)** MEA features of simulations (n=10 per condition) with the ground-truth model parameters (brown), the mode of the posterior (pink), and low probability model parameters (beige). The MEA features are described in Table 2. Data shows mean ± sd. Each feature is normalized by sd_f,prior_, the standard deviation of that feature of simulations sampled from the prior.

As an initial evaluation of SBI, we performed parameter inference on simulated data with known model parameters. The resulting posteriors contained the ground-truth model parameters in a high probability region and the mode of the posterior was close to - or completely overlapped with - the ground-truth (Figure1B, Supplemental Figure 1A). This illustrates that SBI can correctly identify the parameters of our biophysical computational model. Simulations using the mode closely resembled the input data on which the inference had been carried out, while model parameters with low posterior probability generated simulations that were distinctly different from the data (Figure1C,D Supplemental Figure 1B).

### SBI can identify the entire landscape of parameters consistent with MEA measurements of healthy hiPSC-derived neuronal networks

Next, we used SBI to estimate the *in silico* model parameters able to simulate network activity of healthy *in vitro* neuronal networks on MEA. For this, we used 10-minute measurements of spontaneous activity from neuronal networks derived from a healthy individual (C1 in [3]). Simulations with the mode of the posterior distribution were highly similar to experimental measurements, with no significant differences between most MEA features (Figure 2A, Supplemental Figure 2A). To check the reproducibility, we also analyzed measurements from another MEA batch derived from the same individual. Interestingly, posterior distributions looked markedly different for measurements of networks from different MEA batches or with different astrocyte batches (f.e. Figure 2B vs. Supplemental Figure 2B). This suggests that posterior distributions should only be compared when the experimental measurements originate from the same MEA experiment, in line with previous findings [2].

**Figure 2:**
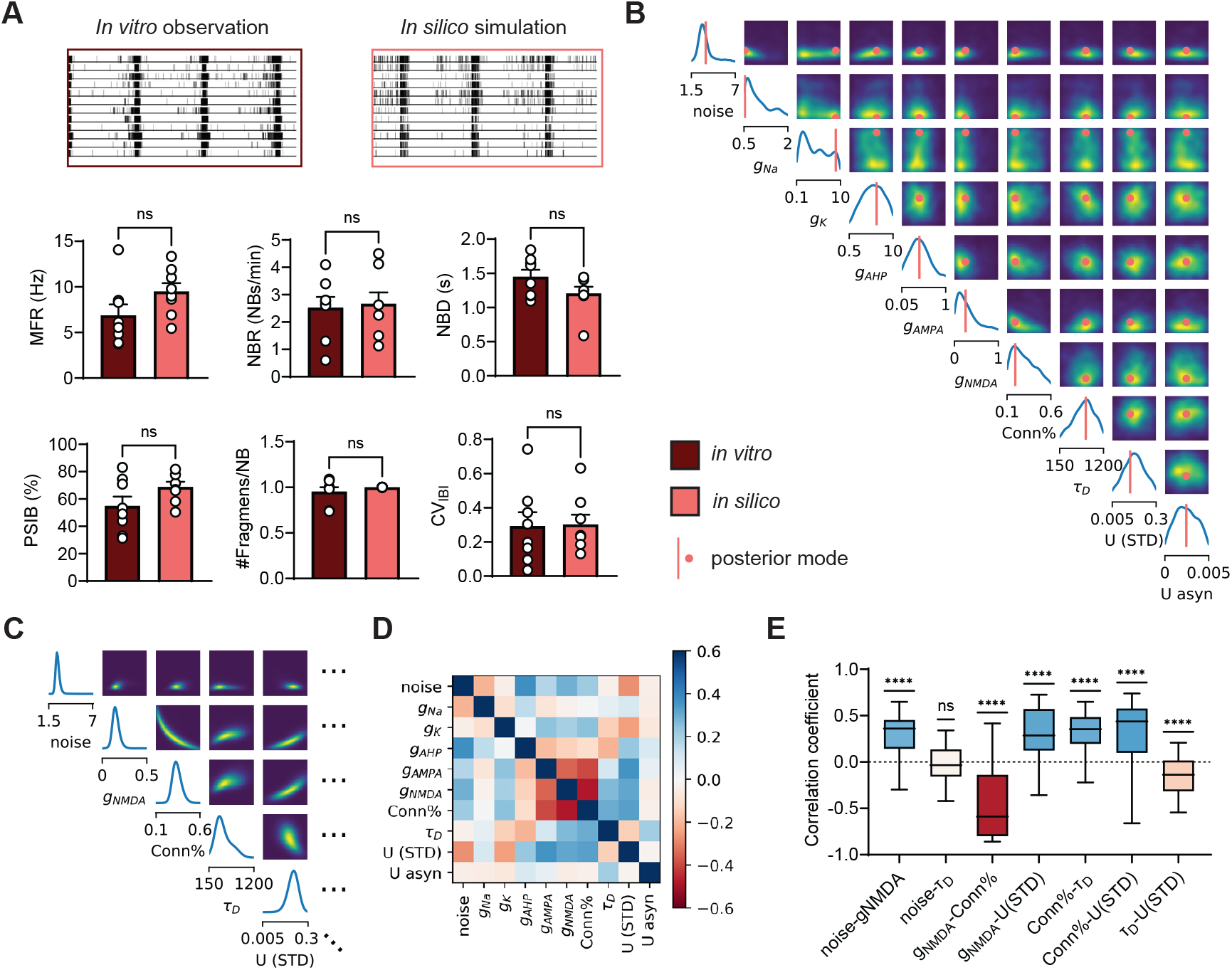
SBI identifies finely-tuned parameters able to generate simulations with high similarity to hiPSC-derived neuronal network measurements. **A)** Representative rasterplot (top left) of spontaneous in vitro neuronal network activity on MEA derived from a healthy individual, used to obtain the posterior distribution, and representative rasterplot (top right) from an *in silico* simulation with parameters from the mode of the inferred posterior distribution. Bottom shows the similarity between features of *in vitro* measurements (n=8) and *in silico* mode simulations (n=8). The features are: mean firing rate (MFR), network burst (NB) rate (NBR), NB duration (NBD), percentage of spikes in NBs (PSIB), the number of fragments per NB (#FBs), and the coefficient of variation of the inter-burst-intervals (CV_IBI_). Data shows mean ± SEM. Data are compared with a Mann-Whitney test where ns means P>0.05. **B)** Inferred posterior distribution. **C)** A subset of a conditional distribution with a particular parameter set. Plots on the diagonal show conditional distribution when all other parameters are fixed. For off-diagonal plots we keep all but two parameters fixed. Distributions are much more narrow compared to the posterior shown in panel B. Note the ranges on the x-axis differ. **D)** Conditional correlation matrix averaged over 50 conditional distributions. **E)** Boxplots of correlation coefficients across 50 conditionals. Colors correspond to average values in panel D. A one-sample t-test was performed to test whether correlation coefficients significantly differed from zero. ns P>0.05, ^****^ P<0.0001.

We noticed that the estimated model parameter distributions, as well as the pairwise marginals, were rather wide, indicating that a large range of different parameters can reproduce the experimental data (Figure 2B). This could indicate one of two possibilities: either changes in some parameters barely affect the network activity, or changes in several parameters do influence network activity, but their effects compensate for each other. To investigate this, we first assessed how sensitive the model is to variations in different parameters by performing a sensitivity analysis on the posterior distribution. We found that although the model is more sensitive to some parameters than others, no sensitivity scores were negligible (Supplemental Figure 2D). Thus, we next investigated whether the wide range of possible parameter combinations could be due to parameters compensating for each other. By examining the pairwise marginals, it seems that parameters are only weakly correlated, as indicated by large clouds without a clear direction (Figure 2B). However, pairwise marginals are averages over many network configurations, where all other model parameters may have various values. Even if certain model parameters can vary widely, each value may require a specific configuration of the remaining model parameters. To explore this further, we held all but one or two parameters constant at values sampled from the posterior distribution and observed what values the remaining model parameters could take to still produce the desired network activity. This resulted in the conditional posterior distribution (Figure 2C, Supplemental Figure 2C). We found that these conditional distributions were significantly narrower, confirming that if some model parameters have a certain value, the remaining model parameters can only vary within a limited range to still yield simulations that match the experimental observation.

When looking at remaining possibilities with all but two parameters set to a constant value (Figure 2C), we observed high correlations between some model parameters. When generating the conditional distribution with many possible configurations, we noticed that some conditional correlations were preserved, leading to an average correlation coefficient significantly different from zero (Figure 2D,E). We found, for example, a strong negative correlation between the AMPA conductance, NMDA conductance, and the number of synapses, indicating that fewer synapses can be compensated by stronger synapses and vice versa. Less intuitively, we found a strong positive correlation between the NMDA conductance and the strength of short-term synaptic depression (U (STD)). Literature suggests that both mechanisms strongly but inversely influence the NB duration (NBD) MEA feature [17, 11]. To investigate this, we computed the sensitivity of the NBD MEA feature to the model parameters and indeed found NBD to be highly sensitive to both NMDA conductance and the strength of STD (Supplemental Figure 2E), suggesting these parameters can compensate for each other. To summarize, we show that SBI can identify model parameters able to replicate neuronal network activity on MEA and that these parameters require fine-tuning with respect to each other, even in the presence of wide univariate marginals.

### SBI can identify pharmacological targets in *in vitro* neuronal networks

To assess whether SBI could identify relevant changes in the physiological properties of hiPSC-derived neuronal networks, we applied it to activity measurements performed on healthy neuronal networks before and after the addition of drugs affecting specific molecular mechanisms. Using SBI, we inferred the posterior distribution for every network before and after the addition of the drug. We compared these distributions per network to see if the parameter differences represented the affected molecular mechanisms.

First, we applied SBI to MEA measurements before and after the addition of Dynasore, which inhibits synaptic vesicle recycling thereby enhancing the amount of STD [18, 19]. In our *in silico* model, the U (STD) parameter governs the amount of neurotransmitters depleted from the readily releasable pool upon every pre-synaptic spike, influencing synaptic depression. The τ_D_ parameter governs the rate of synaptic vesicle recycling: a higher value indicates slower recycling and thus a longer depression of the synapse. We would therefore expect that Dynasore mainly increases τ_D_. When inspecting an example posterior distribution before and after Dynasore addition, it becomes apparent that multiple parameter distributions and their pairwise marginals differ (Figure 3A). Simulations with the mode of both posterior distributions appear highly similar to experimental observations, and the effect of Dynasore on MEA features is very similar in simulations and *in vitro* (Figure 3B). Specifically, the NBD was substantially reduced in both cases. We then compared the univariate marginals before and after the addition of Dynasore per network. For one neuronal network (Figure 3C), we saw a significant shift towards lower values in the marginal of the noise and connectivity, and towards higher values in the marginal of τ_D_ and U. To investigate whether these parameter changes were consistent among many networks, we evaluated the averaged P-values (Figure 3D). We noticed that while the shift in connectivity was large in some networks but not in all (large spread), the shift in the τ_D_ distribution was highly significant in all networks (small average P-value and small spread) (Figure 3D). This demonstrates the importance of comparing multiple networks to eliminate the parameter changes occurring by chance or other factors besides the affected molecular mechanisms.

**Figure 3:**
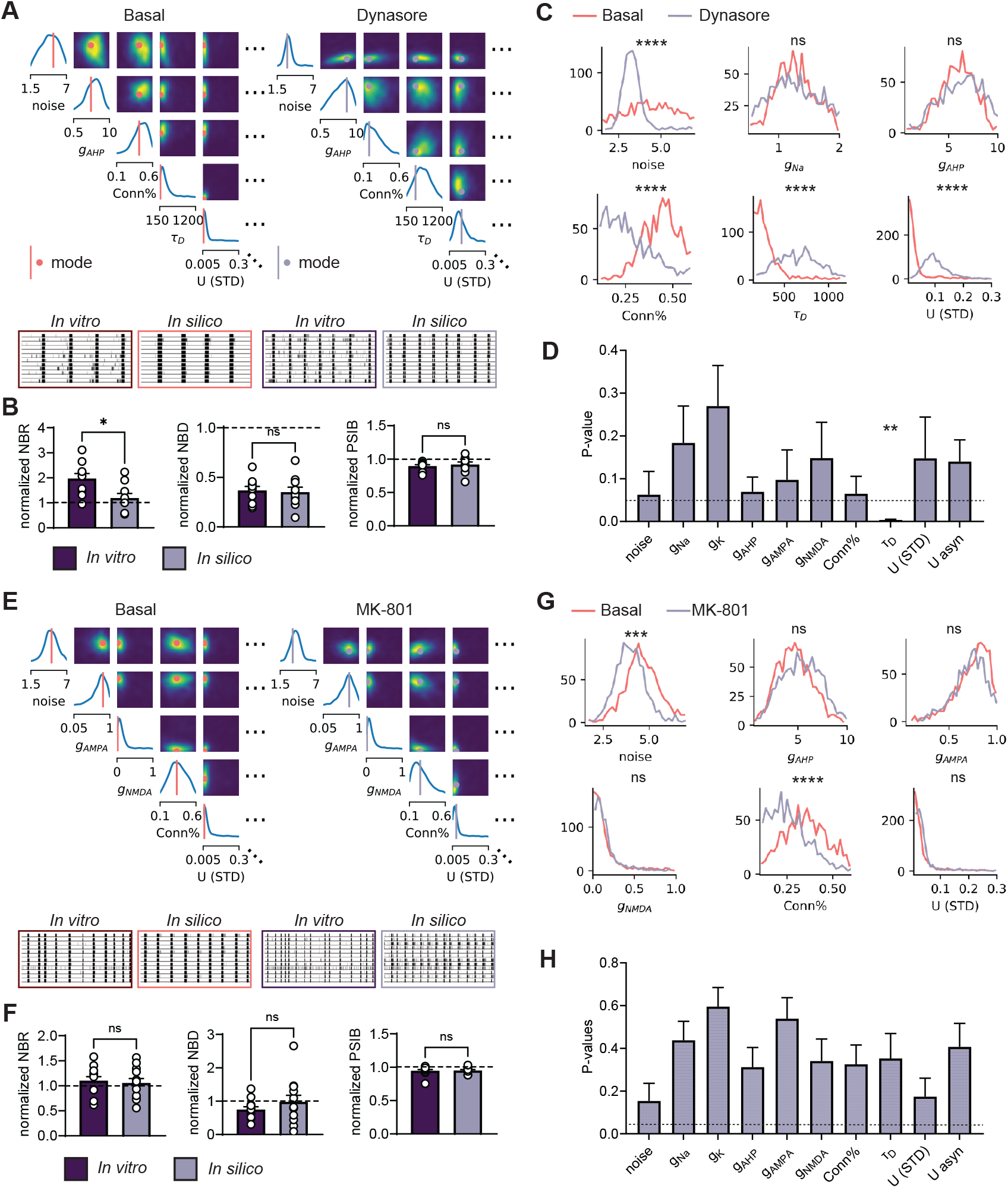
SBI can identify pharmacological targets when network activity is sufficiently altered by drugs. **A)** An example subset of the posterior distributions inferred from a network activity measurement before (left) and after (right) addition of Dynasore (10 µM), and a one-minute rasterplots from the *in vitro* activity and simulated activity with the mode of the posteriors. **B)** Normalized features of experiments (n=10) and simulations (n=10) before and after adding Dynasore. The features are: mean firing rate (MFR), network burst (NB) rate (NBR), NB duration (NBD), and percentage of spikes in NBs (PSIB). Data shows mean ± SEM. Mann-Whitney test was performed between groups. ns P>0.05, ^*^ 0.01>P>0.001. **C)** Univariate marginals from the posteriors shown in panel A. Marginals were compared using a Kolmogorov-Smirnov (KS) test with 50 samples per marginal. ns P>0.05, ^****^ P<0.00001. **D)** Average P-values of the KS-test result of 10 network comparisons per parameter. Data shows mean ± SEM. ^**^ Average P-value<0.01. **E)** An example subset of the posterior distributions inferred from a network activity measurement before (left) and after (right) addition of MK-801 (1 *µ*M), and a one-minute rasterplots from the *in vitro* activity and simulated activity with the mode of the posteriors. **F)** Normalized features of experiments (n=12) and simulations (n=12) before and after adding MK-801. Mann-Whitney test was performed between groups. ns P>0.05. **G)** Univariate marginals from the posteriors shown in panel E. Marginals were compared using a Kolmogorov-Smirnov (KS) test with 50 samples per marginal. ns P>0.05, ^***^ P<0.001, ^****^ P<0.0001. **H)** Average P-values of the KS-test result of 12 network comparisons per parameter.

Next, we applied SBI to MEA measurements before and after the addition of MK-801, an NMDA-receptor blocker [20]. We would expect that MK-801 mainly decreases g_*NMDA*_. However, the posterior distribution before and after the addition of MK-801 revealed minor differences (Figure 3E). Simulations with the mode of the posterior distributions resemble *in vitro* measurements, but the differences between before and after the addition of MK-801 are minimal in both cases (Figure 3F). In some examples, the distributions of the noise and connection parameters significantly shift towards lower values (Figure 3G), but these changes were not consistent across networks (Figure 3H). This suggests that SBI cannot identify molecular changes that do not significantly or consistently affect the neuronal network activity pattern.

### SBI pinpoints known disease mechanisms in patient-derived and genome-edited neuronal networks

To examine whether SBI could identify disease mechanisms in patient-derived neuronal networks, we compiled data from previously investigated and published cell lines. We computed posterior distributions from healthy control networks and compared them to the distributions of patient-derived or genome-edited networks. Because we previously observed that different batches affect SBI results, we only compared posterior distributions of networks cultured and measured on the same MEA plate.

First, we analyzed data from neuronal networks derived from patients with Dravet Syndrome (DS) and Generalized Epilepsy with Febrile Seizures plus (GEFS+), both epileptic encephalopathies linked to a variant in SCN1A, which encodes part of the voltage-gated sodium channel Na_V_1.1 (PAT001 GEFS and PAT001 DS from [7]). These networks were cultured alongside networks derived from a healthy individual (Control). Previous *in vitro* experiments revealed lower sodium current densities, altered action potential (AP) decay dynamics and firing patterns suggestive of a reduction in potassium currents, and reduced spontaneous excitatory post-synaptic current (sEPSC) frequency and amplitudes in GEFS+ and DS networks compared to Control [7, 10]. Moreover, increased STD was recently suggested to be important in GEFS+ and DS networks to explain the phenotype [11]. We found that the posterior distributions inferred with SBI visually differed between healthy control networks and DS and GEFS+ (Figure 4A). Univariate marginals of several parameters also significantly differed in both independent MEA batches (Figure 4B and Supplemental Figure 3A). Consistent over two batches, the noise, sodium conductance, and potassium conductance distributions significantly shifted towards lower values in both GEFS+ and DS (Figure 4C). The strength of STD distribution significantly shifted towards higher values. Additionally, DS-inferred distributions showed a shift towards higher NMDA conductances, and GEFS+-inferred distributions showed a shift towards lower connectivity and higher asynchronous release.

**Figure 4:**
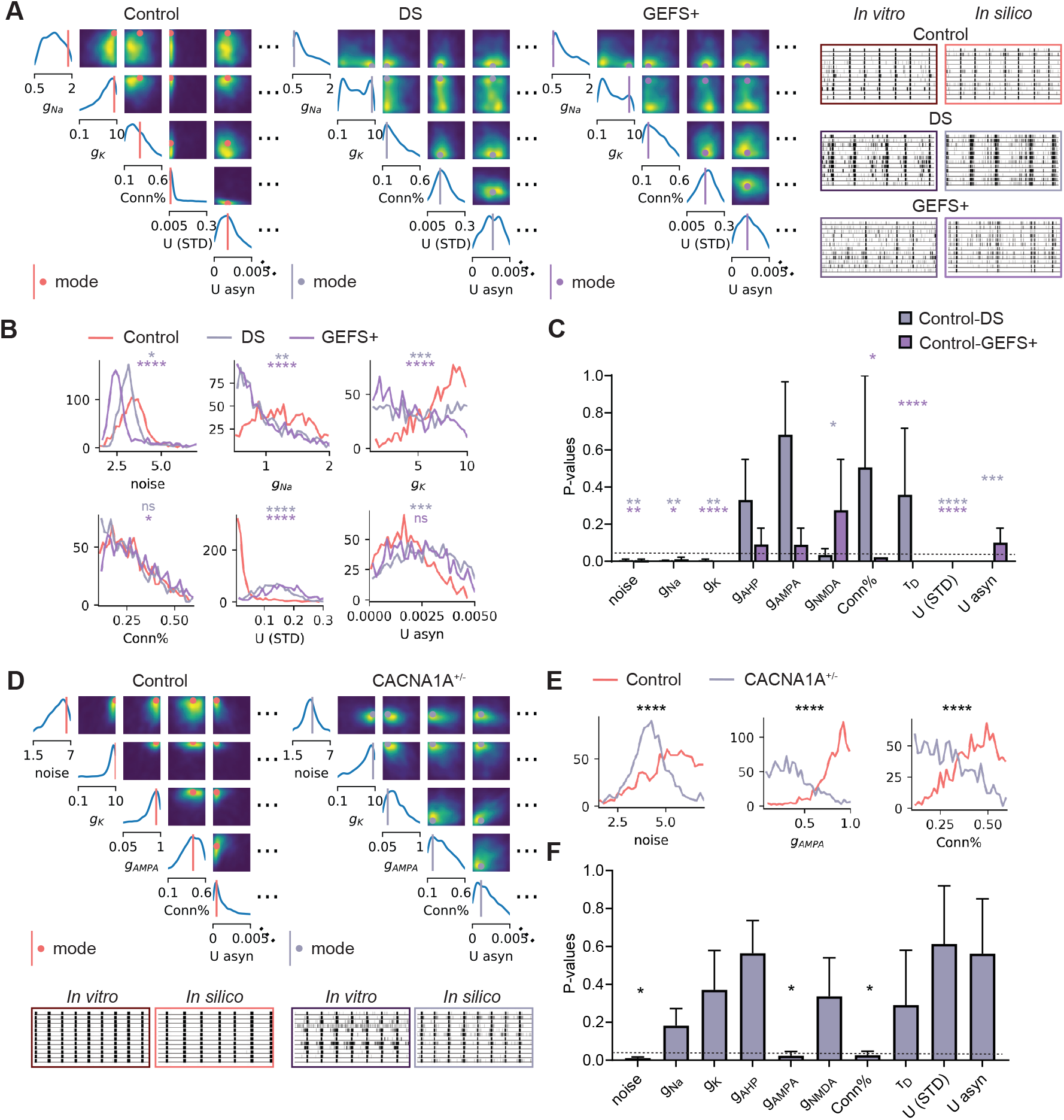
SBI identifies known disease mechanisms in patient-derived and genome-edited neuronal networks. **A)** Left: Example subsets of posterior distributions inferred from average neuronal network activity of a healthy Control, a patient with Dravet Syndrome (DS), and a patient with Generalized Epilepsy with Febrile Seizures plus (GEFS+). Right: representative 1-minute rasterplots of spontaneous network activity of the different cell lines and the corresponding simulations with the mode of the posterior distributions. **B)** Univariate marginals from the posteriors shown in panel A. Marginals were compared using a Kolmogorov-Smirnov (KS) test with 50 samples per marginal. ns P>0.05, ^*^ P<0.05, ^**^ P<0.01, ^***^ P<0.001, ^****^ P<0.0001. **C)** Average P-values of the KS-test result of 2 batches per parameter. Data shows mean ± SEM. Average P>0.05, ^*^ Average P<0.05, ^**^ Average P<0.01, ^***^ Average P<0.001, ^****^ Average P<0.0001. **D)** Top: Example subsets of posterior distributions inferred from average neuronal network activity of Control and CACNA1A^+/−^ networks. Bottom: representative 1-minute rasterplots of the spontaneous network activity from the isogenic lines and corresponding simulations with the mode of the posterior distributions. **E)** Univariate marginals from the posteriors shown in panel D. **F)** Average P-values of the KS-test result of 3 batches per parameter.

Second, we analyzed networks differentiated from isogenic iPSC lines with a monoallelic frameshift variant in exon 8 of *CACNA1A*, generated via CRISPR/Cas9 [21]. *CACNA1A* encodes part of the voltage-gated calcium channel Ca_V_1.1, crucial for proper neurotransmitter release [22]. These networks were cultured alongside with networks derived from a healthy individual (Control). *in vitro* experiments revealed a significantly reduced number of synapses, altered AMPA signaling, and reduced potassium channel functioning. Per MEA, we analyzed the shift in univariate marginals between the isogenic networks (Figure 4E). Averaged over three batches (Figure 4F, Supplemental Figure 3B), we found a significant shift towards lower values in the distributions of the noise parameter, the AMPA conductance, and the connectivity. There was a significant negative shift in the potassium conductance in only one batch.

## Discussion

Utilizing hiPSC-derived neuronal networks on MEA, whether genome-edited or patient-derived, provides a rapid means to gather neuronal electrophysiological data. These data often exhibit distinct characteristics that differ between healthy and diseased networks, reflecting underlying pathological molecular mechanisms [2, 3, 5, 6, 7, 21]. While some mechanisms are explicitly linked to specific activity patterns [17, 11], characterizing them definitively through quantitative analysis of the data remains challenging. Additional experiments to nonetheless uncover these mechanisms, such as patch-clamp measurements or RNA-sequencing, are costly and time-consuming and may not cover all possible molecular pathways, potentially overlooking crucial mechanisms [10, 21]. Here, we propose the use of simulation-based inference (SBI) and our previously developed biophysical computational model to automatically identify disease mechanisms in cultured neuronal networks on MEAs. While SBI shows promise for accurately inferring biological and biophysical parameters [23], its application to uncover pathophysiology in MEA data has not been previously reported.

To investigate whether SBI could pinpoint specific molecular pathways affected in neuronal networks, we applied it to pharmacologically modulated networks. Using Dynasore, an endocytosis inhibitor [18], SBI accurately predicted an increase in the time constant governing synaptic vesicle endocytosis across all networks. Additionally, in most networks, SBI predicted a reduction in network connectivity and an increase of U, describing the strength of STD. This aligns with the expected outcome, as prolonged synaptic vesicle depletion leads to enhanced synaptic depression and reduced functional connectivity [19]. Using MK-801, an NMDAr-blocker, SBI predicted a decrease in connectivity, but this prediction was not consistent among all networks. Moreover, SBI predicted an initial NMDAr conductance close to zero. This appears in line with the lack of a clear effect of MK-801 on the MEA activity features in most networks. This shows that SBI can only detect neuronal network changes that are clearly and consistently reflected by a change in MEA features describing the network behavior.

Using cultured neuronal networks from patients with GEFS+ and DS, both harboring a variant in *SCN1A* encoding part of the voltage-gated sodium channel, SBI successfully predicted lower values of the voltage-gated sodium channel conductance for both diseases, with the change more pronounced in DS across the two batches. This is in agreement with voltage clamp measurements in GEFS+ and DS neurons, where DS neurons show more severely impaired sodium currents [7]. SBI also predicted lower values of the voltage-gated potassium channel conductance in both GEFS+ and DS, albeit more prominently in GEFS+ on average. This appears in line with the current clamp measurements demonstrating impaired AP decay dynamics in both conditions, albeit more markedly in GEFS+ [7]. Additionally, SBI forecasted a significant increase in STD strength in both GEFS+ and DS, along with a larger STD time constant for DS and increased asynchronous release in GEFS+. These predictions align with previous findings implicating these mechanisms in the observed phenotypes of GEFS+ and DS [11]. Furthermore, lower sEPSC amplitudes and frequencies were observed in DS neurons, possibly indicating more depressed synapses [10]. No changes in the number of synapses were found between healthy and DS neurons, which is also predicted by SBI [10]. This was not investigated for the GEFS+ neurons.

Next, we investigated CRISPR/Cas9 edited networks with haploinsufficiency of *CACNA1A*, a gene encoding part of the voltage-gated calcium channel. Averaged across three independent batches, SBI predicted a significantly lower connection probability in *CACNA1A*^+/−^ neuronal networks. This aligns with the immunocytochemistry results showing a significantly reduced number of synapses in CACNA1A^+/−^ networks [21]. Furthermore, SBI suggested a reduction in AMPA conductance. Although direct measurements of the AMPA current were not conducted *in vitro*, pharmacological testing revealed *CACNA1A*^+/−^ network activity was governed more by GluA2-lacking AMPA receptors. Our computational model only contains GluA2-containing AMPA receptors with fast opening and closing dynamics and NMDA receptors with slower opening and closing dynamics and dependence on repetitive firing. GluA2-lacking AMPA receptors resemble the dynamic of the NMDA receptors present in our *in silico* model [24]. Therefore, a shift towards more influence of GluA2-lacking AMPA receptors *in vitro*, could be predicted by SBI as a reduction in GluA2-containing receptors, leading to a more NMDA-dominated synaptic transmission. In *CACNA1A*^+/−^ neurons, evidence was also found for reduced voltage-gated potassium channel function, indicated by the slightly lower AP decay time, reduced AHP amplitude, reduced rheobase, and altered potassium channel-related RNA expression [21]. SBI predicted a significant shift towards lower values of the potassium-channel conductance. However, this shift was observed in only one batch, reducing the overall significance of the effect when averaged across three batches. The lack of an effect in the other batches might be due to the small size of the effect measured *in vitro*, or to the model’s relative insensitivity to the potassium channel parameter (Supplemental Figure S2D).

SBI predicted a lower amplitude of the noisy membrane potential fluctuations in GEFS+, DS, and *CACNA1A*^+/−^ networks. This noise variable represents both the effect of spontaneous neurotransmitter release on the membrane potential - which is dependent on the number of synapses, their strengths, and their spontaneous release frequency - and the effect of electrical noise on the membrane potential - which is dependent on the input resistance of the neurons, their size, and the propensity of ion channels to randomly open and close [25]. In DS neurons, we recorded lower sEPSC amplitudes and frequencies, which would explain the decreased effect of these events on the membrane potential fluctuations. In *CACNA1A*^+/−^ neurons, the synaptic calcium channels are impaired. Spontaneous opening of these channels underlies part of the spontaneous neurotransmitter release, and blocking these channels has been shown to reduce miniature EPSC frequency [26]. Thus, the impaired noise in *CACNA1A*^+/−^ neurons may be explained by the impaired calcium channels. However, *CACNA1A*^+/−^ neurons also showed significantly increased soma sizes, which would decrease the input resistance and thus reduce the effect of channel noise on the membrane potential, which may also explain a reduction in the noise parameter [25].

### Limitations and advantages

Employing SBI with our biophysical model presents several challenges. First, we assumed that our model included all relevant biological processes to, in principle, simulate the experimental observations. However, we cannot exclude that certain disease mechanisms may not be fully captured, due to inherent limitations of our model. For instance, pathologies arising from inhibitory neurons [27] or mitochondria [6], may not be faithfully simulated. Second, our model includes a large number of parameters. We decided to infer ten parameters, keeping the other parameters constant. We argued that these represent the most important neuron and synapse properties. However, not all experimental measurements could be perfectly replicated by the model, as illustrated in Figure 3 and Supplemental Figure S2. This may have resulted from the inability of the model to vary the remaining parameters. At the same time, the addition of more free model parameters hampers the interpretability of the resulting posterior distribution. Already with ten model parameters, the univariate marginals are wide, and it is difficult to interpret the model parameters in a high dimensional space. More model parameters will likely widen the marginals further, as more model parameters can compensate for each other, reducing its sensitivity to detect changes between healthy and diseased networks. Adding more model parameters also requires more simulations to train the NDE, which is computationally expensive despite the suitability of SBI to work with high-dimensional parameter spaces [23].

Another limitation of SBI is the need to define prior distributions and features describing the experimental data, which can introduce biases. We defined a prior distribution based on biological plausibility as well as experience with model parameter configurations resulting in realistic MEA feature values. We defined MEA features often used in studies of neuronal networks on MEA [2], supplemented with MEA features describing the other characteristics of the data. Many of these MEA features depend on the proper detection of NBs, which can be difficult in the largely varying set of simulations [28]. To limit biases, NB detection and MEA feature calculation have been performed identically in the *in vitro* measurements and *in silico* simulations. Nevertheless, some differences in NB detection, for instance resulting from different signal-to-noise ratios, could explain some discrepancies between simulations and experimental observations.

A final limitation of SBI is the difficulty of verifying the correctness of the inference. We performed posterior predictive checks to see if the ground-truth parameters were contained within high-probability regions, but we lacked the means to validate if the entire posterior was estimated properly. Since the likelihood of our model is intractable, computing a ground-truth posterior distribution for comparison was not feasible. Consequently, corroborating SBI predictions with targeted *in vitro* experiments is imperative when investigating disease mechanisms.

Despite these limitations, we demonstrate that SBI offers several advantages over traditional parameter estimation methods. While trial-and-error methods are common in neuronal modeling studies, they lack systematicity and often yield singular solutions [10, 29, 30]. Similarly, parameter-searching methods, such as grid searches or evolutionary algorithms, require defining distance measures to experimental observations, necessitate numerous simulations for every experimental observation, and provide only one optimal parameter set without quantifying uncertainty or parameter importance [12, 13]. Furthermore, numerous studies highlight parameter degeneracy in biological neuronal networks and computational models: the same activity can arise from vastly different parameters [31, 14]. Recognizing this *nonuniqueness* is crucial to avoid drawing incomplete conclusions when observing changes in parameter values between experimental conditions. SBI addresses these challenges by identifying the full range of parameters that explain the observations, unlike other parameter estimation techniques. Finally, SBI is amortized, meaning that performing the many simulations and training the NDE only need to be executed once. If new experimental data becomes available, a corresponding posterior distribution can be calculated in a matter of seconds. Therefore, posterior analysis could be a plausible addition to the standard MEA data analysis pipeline.

### Recommendations and Conclusion

Based on our findings, we propose that SBI holds promise for automatically detecting disease mechanisms in patient-derived neuronal networks on MEA. However, several important recommendations and considerations should be taken into account when employing this method. Firstly, it is crucial to compare neuronal networks cultured on the same MEA and measured simultaneously, as differences in astrocyte and MEA batches can introduce confounding factors. Principal-components analysis conducted previously highlighted the influence of these factors on MEA data features [2], and here we observed their effect on SBI-identified posterior distributions. Secondly, conducting multiple comparisons using independent neuronal preparations on MEA is essential to discern changes consistent across batches. Previous advice emphasized the importance of employing sufficient experimental batches [2], and here we observed that not all parameter differences were reproduced across batches. We thus suggest comparing univariate marginals per MEA and averaging resulting P-values across all MEAs to identify parameters that are altered consistently. Thirdly, targeted *in vitro* investigations of SBI-predicted disease mechanisms are imperative to validate predictions. Besides posterior-predictive checks, it is not possible to check the correctness of the inference performed by SBI. Additionally, SBI identifies a multitude of parameter possibilities consistent with the data, resulting in multiple predicted parameter changes of which potentially only a subset is present *in vitro*. Investigating all possibilities suggested by SBI can provide clarity on genuine *in vitro* changes. Nevertheless, SBI significantly reduces the number of *in vitro* experiments required to properly identify disease mechanisms. Without SBI, researchers would face the daunting task of conducting endless experiments to explore all possibilities, or risk overlooking crucial mechanisms by basing experiments solely on initial hypotheses. With SBI, a trained NDE, and the analysis pipeline outlined here, MEA activity can be analyzed easily and rapidly to automatically identify all the possible disease mechanisms able to explain patient-derived neuronal network phenotypes. This paves the way for targeted experiments and novel insights into disease.

## Materials and Methods

### *in vitro* MEA data acquisition

We used data from neuronal networks derived from hiPSCs, previously cultured and recorded on MEA. hiPSC cells were differentiated into excitatory cortical Layer 2/3 neurons through doxycycline-inducible overexpression of Neurogenin 2 (*Ngn2*) as described previously [1, 2].

The control line used for both pharmacological experiments and as control with diseased networks was derived from a 30-year-old male, as previously described and characterized [3, 32].

Drug manipulations were previously performed on neuronal networks derived from the control line as described in [11] for Dynasore and in [10] for MK-801, using the 24-well MEA system (Multi-channel systems, MCS GmbH, Reutlingen, Germany), with a sampling frequency of 10 kHz.

MEA recordings of neuronal networks derived from patients with GEFS+ and DS were kindly provided by Nael Nadif Kasri and Eline van Hugte, and are described (Control, PAT001 GEFS, PAT001 DS) in [7]. Activity was recorded on the 24-well MEA system (Multichannel systems, MCS GmbH, Reutlingen, Germany) for 10 minutes with a sampling frequency of 10 kHz.

MEA recordings of control and CACNA1A^+/−^ networks were kindly provided by Nael Nadif Kasri and Marina Hommersom, and are described in [21]. Activity was recorded on the Maestro Pro MEA system (Axion BioSystems, Atlanta, GA, USA) for 5 minutes with a sampling frequency of 12.5 kHz. Because this MEA system had 16 electrodes per well, we omitted the corner four electrodes from analysis, to mimic the electrode topology of the Multichannel MEA system and the computational model.

During all recordings, the temperature was maintained at 37 ° C, and a flow of humidified gas (5% CO_2_ and 95% ambient air) was applied onto the MEA plate.

### Computational Model

The *in silico* computational model is described in [10], with the addition of asynchronous neurotransmitter release described in [11]. In short, the model consists of one hundred Hodgkin-Huxley neurons with leaky channels and voltage-gated sodium and potassium channels. We also include sAHP channels, modeled as potassium channels whose conductance increases upon AP firing. The neurons are heterogeneously excitable through a variable constant external input current, accounting for intrinsic differences. In addition, we induce stochastic fluctuations of the membrane potential of neurons to mimic the effect of spontaneous neurotransmitter release or channel noise. Neurons are connected to a fraction of other neurons through synapses. These synapses include models of AMPA receptors (AMPArs), which open immediately upon arrival of a pre-synaptic spike and close rapidly. Additionally, we include NMDA receptors (NMDArs), which open and close slowly. The NMDArs are blocked by magnesium ions that are removed upon depolarization of the post-synaptic neuron. The strengths of the synapses vary slightly and are modulated by STD, following the Markram-Tsodyks model [33].

Pre-synaptically, neurotransmitters are released synchronously (time-locked to the arrival of the AP), and asynchronously (where the release probability increases upon repetitive firing), following the Markram-Tsodyks model extension by Wang et al. [34]. Fixed parameter values and prior ranges of the free parameters can be found in Table 1. Neurons were placed on a grid, allowing for the inclusion of distance-dependent conduction delays and virtual electrodes, mimicking MEA measurements.

Forward simulations were performed using the brian2 simulator [35] on a dedicated platform with synaptic parallel computing for dynamical load balancing also used in [36].

### Pre-processing and feature extraction

*in vitro* MEA recordings and *in silico* virtual electrode recordings are pre-processed and analyzed identically. Signals were filtered between 100 and 3500 Hz using a fifth-order Butterworth filter. We detected spikes using an amplitude threshold-based method, where the threshold was four times the root mean square of the electrode signal.

The computed MEA features as summarized in Table 2. The network firing rate was computed by binning spikes at all electrodes in 25 ms time bins. To detect NBs, we employed two thresholds on this network firing rate to start and stop the NB, set to 1/4th and 1/50th of the maximum firing rate respectively. Additionally, 50% of the active electrodes (i.e., electrodes with an average firing rate above 0.02 Hz) had to be firing during the NB, and the firing rate should remain above the NB start-or stop-threshold for 50 ms to start or end NB detection. To detect NB Fragments, we smoothed the network firing rate by convolution with a Gaussian kernel and used a peak detection algorithm on the smoothed network firing rate where peaks should have a minimal height of 1/16th of the maximum firing rate and a minimal prominence of 1/20th of the maximum firing rate. From this NB detection, we constructed five features. Specifically, we calculated the NB Rate (NBR), describing the number of NBs per minute, the NB Duration (NBD), describing the time between the start and stop of the NB in seconds, the Percentage of Spikes in NBs (PSIB), describing the percentage of spikes contained in the detected NBs, the number of fragments per NB (#FBs), and the coefficient of variation of the inter-NB-intervals (CV_IBI_).

We computed eight additional features describing other characteristics of the data. The first feature was the mean firing rate (MFR). Then we computed the continuous inter-spike-interval (ISI) time-series per MEA electrode, by determining the time between the previous and next detected AP for every time point. From these ISI-time-series, we computed the average ISI (mean ISI) across all timepoints and electrodes. By first taking the average ISI-time-series across electrodes, we defined the standard deviation of the ISI over time (sd ISI temp), and by first taking the average ISI across time points, we defined the standard deviation of the ISI over electrodes (sd ISI elec). Moreover, we computed the sample correlation coefficient between every pair of electrodes ISI-time-series. From this, we computed the average correlation coefficient (mean CC) and the standard deviation of the correlation coefficients (sd CC). Finally, we computed the average ISI-distance between every pair of electrodes as described in [37]. The above measures describe the spiking behavior of a network as well as the variation of spiking behavior temporally and spatially. Finally, to have a measure for the strength of oscillations in the network activity without relying on NB detection, we computed the Maximum Autocorrelation Component (MAC), as described in [38].

We trained the NDE on two additional features, the average and standard deviation of the correlation coefficient between binarized spike trains. However, we observed high correlations (0.96) between these features and the mean CC and sd CC of the ISI-time-series. Consequently, we chose not to show these features in the figures and we argue they could be omitted from training of the NDE.

### Simulation-based-inference

To estimate the posterior distribution of the free parameters of the computational model, we employed amortized single-round neural posterior estimation [23] using the SBI toolbox [16] in a Python 3.9 environment. We sampled 300,000 parameter configurations from the prior distribution (a uniform distribution within the ranges shown in Table 1) and performed simulations with the computational model with each of these configurations. For every simulation, we computed the 13 MEA features described above. Next, we trained an NDE on these parameter configurations and resulting MEA features. This NDE was the standard Mixture Density Network provided by the SBI toolbox. Then, we evaluated the NDE with MEA features from either simulated network activity (for posterior-predictive checks) or experimental network activity. This resulted in a posterior distribution for each observation. Posterior distributions were visualized by plotting the marginals of 1000 samples drawn from the distribution.

### Posterior-predictive checks

As a check for the accuracy of the inference, we performed posterior-predictive checks. Initially, we used ground-truth model parameters to generate simulations. The trained NDE was then used to estimate the posterior distribution of model parameters consistent with these simulations. The univariate and pairwise marginals were then inspected to see whether the ground-truth model parameters fell within regions with a probability above 50% of the maximal probability. We performed posterior-predictive checks with NDEs trained on 100,000, 200,000, and 300,000 simulations, to see when the posterior distributions converged and checks were sufficient (all twenty checks in 50% or higher regions). To visualize the accuracy of the estimation, we performed ten simulations with the mode of the posterior distribution, the ground-truth model parameters, and low-probability model parameters, and quantified the MEA features. For visualization, we normalized the MEA features by the standard deviation of that feature in the 300,000 simulation dataset.

### Comparison to *in vitro* data

We compared MEA features of simulations generated with model parameters from the mode of the posterior distribution to MEA features of the corresponding experimental measurements. We performed statistical analysis using GraphPad Prism 5 (GraphPad Software, Inc., CA, USA). We first assessed the normality of the distributions using a Kolmogorov-Smirnov test. Given that normality of the features was often not ensured, we used non-parametric testing using a Mann-Whitney test to determine whether simulation and experimental features showed significant differences.

### Conditional correlations

To investigate possible compensation mechanisms between model parameters, we generated 50 conditional distributions by holding all but two model parameters constant at values sampled from the posterior distribution and finding the distribution of the remaining pair of model parameters. We then took 50 points from the resulting distribution and computed the Pearson correlation coefficient between the two model parameters, repeating this for every pair of model parameters. This resulted in 50 correlation coefficients per model parameter pair. To assess whether the average correlation coefficient per model parameter pair significantly differed from zero, we performed a one-sample t-test using GraphPad Prism 5.

### Identification of affected parameters

To identify which parameter distributions were affected by drugs or disease, we compared the univariate marginals. First, we calculated the marginals using the MEA features of every network before and after drug application. We then took 50 samples from the marginals from each test and assessed whether the samples originated from different distributions using a Kolmogorov-Smirnov test using the Scipy stats package within a Python environment. This resulted in one P-value per network which we averaged over all networks to get an overall P-value. For comparison between healthy and diseased networks, we averaged the MEA features per cell line and calculated the marginals. Using 50 samples from each set of marginals, we performed a Kolmogorov-Smirnov test, resulting in a P-value per MEA. Finally, we averaged the P-value across all available MEAs.

## Code Availability

The trained neural density estimator (NDE), the training dataset, the code to analyze experimental data and create the posterior distribution, the code to run simulations with the computational model, and the code to create the figures are available on Gitlab (https://gitlab.utwente.nl/m7706783/SBI_MEA_model).

## Acknowledgements

This work was supported by the Netherlands Organisation for Health Research and Development ZonMW; BRAINMODEL PSIDER program 10250022110003 (to M.F.). We thank Maurice van Putten, PhD, for his invaluable support, expertise, and generous provision of the code to implement synaptic parallel computing for dynamic load balancing. Moreover, we thank Eline van Hugte, Marina Hommersom, and Nael Nadif Kasri for providing MEA recordings from patient-derived and genome-edited *in vitro* neuronal networks.

