## Supplemental Figures for "Automated inference of disease mechanisms in patient-hiPSC-derived neuronal networks"

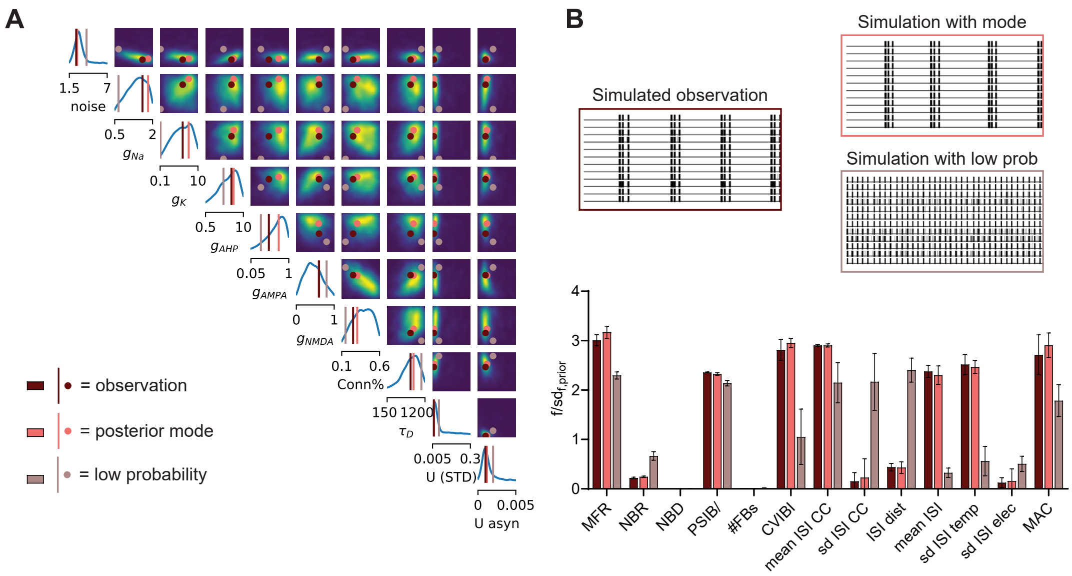


Figure S1: **Posterior predictive check shows SBI can correctly identify ground-truth parameters. A)** Inferred posterior for 10 model parameters given 15 MEA features of simulated activity. Ground-truth model parameters are shown in brown, posterior mode in pink, and low-probability model parameters in beige. **B)** Top: Rasterplots showing 1 minute of (left) simulation used as input for the inference, (top right) example simulation with the mode of the posterior distribution, and (bottom right) example simulation with the low probability model parameters. Bottom: MEA features of simulations (n=10 per condition) with the ground-truth model parameters (brown), the mode of the posterior (pink) and low probability model parameters (beige). The MEA features are: mean firing rate (MFR), network burst (NB) rate (NBR), NB duration (NBD), percentage of spikes in NBs (PSIB), the number of fragments per NB (#FBs), the coefficient of variation of the inter-burst-intervals ($CV_{IBI}$), the average correlation coefficient between channel signals (mean CC), the standard deviation (sd) of the CCs (sd CC), the inter-spike interval (ISI) distance (ISI dist), the mean ISI, the sd of ISIs over time (sd ISI temp), the sd of ISIs between electrodes (sd ISI elec) and the maximum autocorrelaction component (MAC). Data shows mean ± sd.

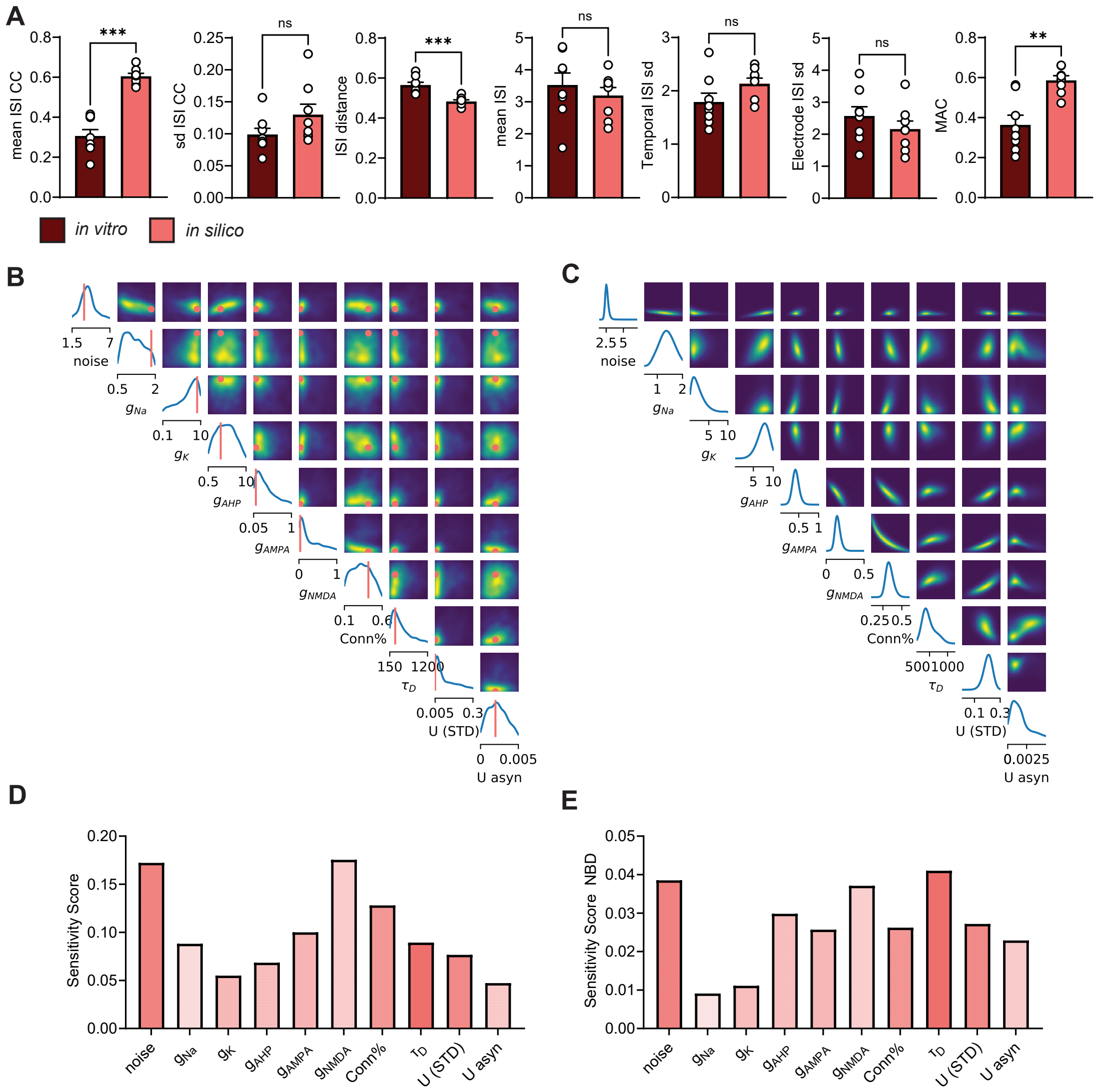


Figure S2: **SBI can correctly identify constrained parameters resulting in simulations mimicking healthy neuronal network behavior and their corresponding sensitivities. A)** MEA features of *in vitro* measurements and simulations with the mode of the posterior (pink). The features are: mean firing rate (MFR), network burst (NB) rate (NBR), NB duration (NBD), percentage of spikes in NBs (PSIB), the number of fragments per NB (#FBs), the coefficient of variation of the inter-burst-intervals ($CV_{IBI}$), the average correlation coefficient between channel signals (mean CC), the standard deviation (sd) of the CCs (sd CC), the inter-spike interval (ISI) distance (ISI dist), the mean ISI, the sd of ISIs over time (sd ISI temp), the sd of ISIs between electrodes (sd ISI elec) and the maximum autocorrelaction component (MAC). Data shows mean ± SEM. **B)** Inferred posterior distribution of healthy neuronal networks of a different MEA batch compared to Figure **??**B (Control in [1]. **C** Complete conditional distribution of the subset shown in Figure **??**C). Plots on the diagonal show conditional distribution when all other parameters are fixed. For off-diagonal plots we keep all but two parameters fixed. **D)** Sensitivity scores computed using eigendecomposition of the posterior shown in Figure **??**B). **E)** Sensitivity scores of the posterior computed with a neural network trained to predict the Network Burst Duration (NBD) feature from the parameters. Scores indicate how much the NBD feature is influenced by the model parameters.

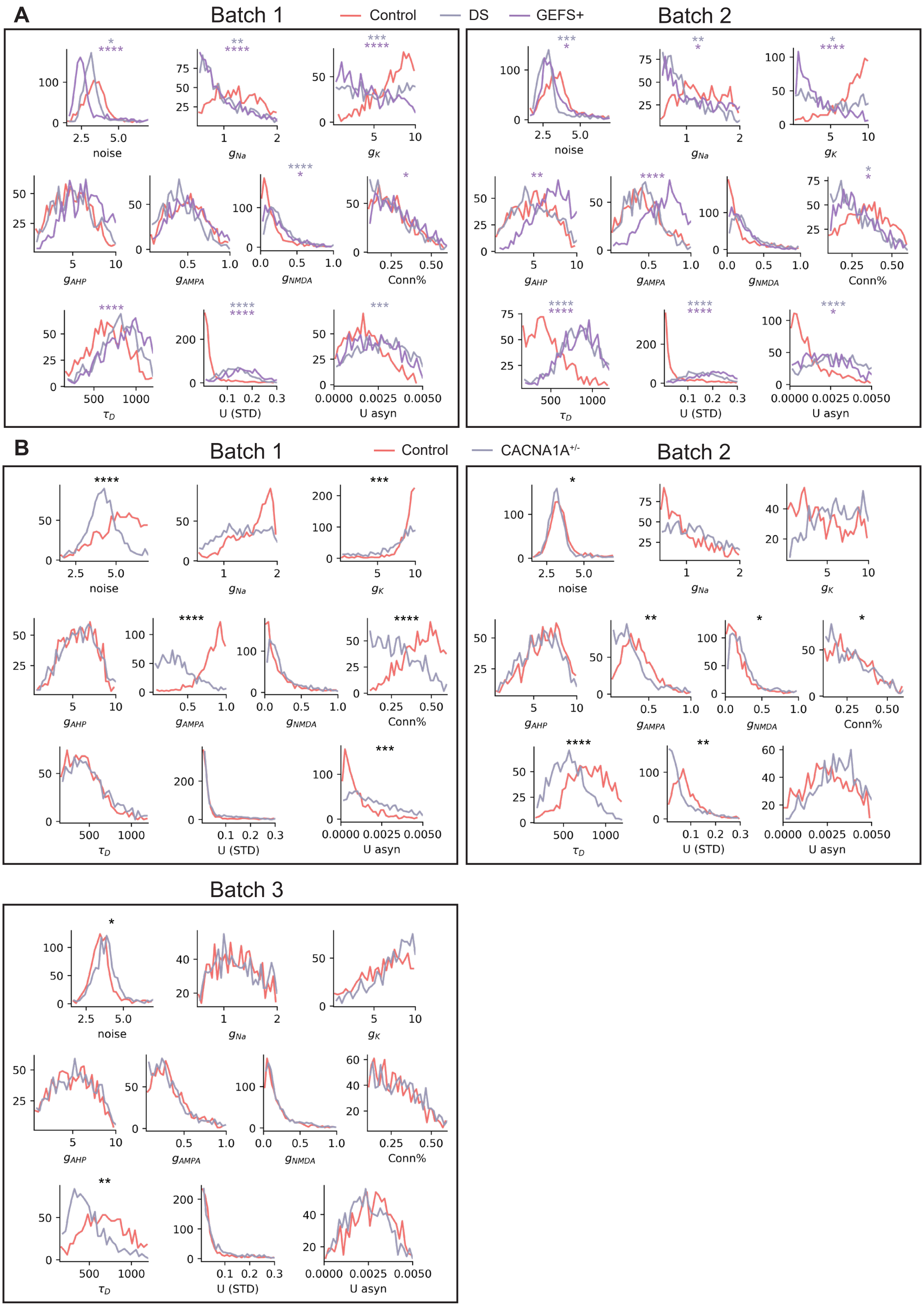


Figure S3: **Comparisons between univariate marginals of healthy and diseased neuronal networks from different batches.** **A)** Comparisons of two batches between neuronal networks of a healthy control, a patient with DS and a patient with GEFS+. **B)** Comparisons of three batches between neuronal networks of a healthy control and CACNA1A deficient networks. Marginals were compared using a Kolmogorov-Smirnov (KS) test test with 50 samples per marginal. * P<0.05, ** P<0.01, *** P<0.001, **** P<0.0001.
